# EB-SUN, a New Microtubule Plus-End Tracking Protein in *Drosophila*

**DOI:** 10.1101/2024.09.11.612465

**Authors:** Sun K. Kim, Stephen L. Rogers, Wen Lu, Brad S. Lee, Vladimir I. Gelfand

## Abstract

Microtubule (MT) regulation is essential for oocyte development. In *Drosophila*, MT stability, polarity, abundance, and orientation undergo dynamic changes across developmental stages. In our effort to identify novel microtubule-associated proteins (MAPs) that regulate MTs in the *Drosophila* ovary, we identified a previously uncharacterized gene, CG18190, encoding a novel MT end-binding (EB) protein, which we propose to name EB-SUN. We show that EB-SUN colocalizes with EB1 at growing microtubule plus-ends in *Drosophila* S2 cells. Tissue-specific and developmental expression profiles from Paralog Explorer reveal that EB-SUN is predominantly expressed in the ovary and early embryos, while EB1 is ubiquitously expressed. Furthermore, as early as oocyte determination, EB-SUN comets are highly concentrated in oocytes during oogenesis. EB-SUN knockout (KO) results in a decrease in MT density at the onset of mid-oogenesis (Stage 7) and delays oocyte growth during late mid-oogenesis (Stage 9). Combining EB-SUN KO with EB1 knockdown (KD) in germ cells significantly further reduced MT density at Stage 7. Notably, all eggs from EB-SUN KO/EB1 KD females fail to hatch, unlike single gene depletion, suggesting a functional redundancy between these two EB proteins during embryogenesis. Our findings indicate that EB-SUN and EB1 play distinct roles during early embryogenesis.

**Significance Statement:**

- Our study identified a novel microtubule end-binding (EB) protein, which is highly expressed in the *Drosophila* ovary, and we propose to name it EB-SUN.
- We demonstrate the functional redundancy of EB-SUN and EB1 during oocyte development, while highlighting their distinct roles in early embryogenesis.
- Given the high conservation of MT-dependent mechanisms in oogenesis and EB proteins across species, these findings provide insights into the differential regulation and tissue-specific functions of the EB family, using *Drosophila* as a model system, potentially benefiting future research on MT regulation in higher organisms.

## Introduction

Understanding the molecular basis of oocyte development is critical due to its direct impact on fertility and reproductive success. The spatiotemporal regulation of microtubule (MT) dynamics is essential for oocyte development. The MT-dependent mechanisms during oogenesis and structural aspects of the germline cyst where oogenesis occurs are highly conserved across species (Lei & Spradling, 2016; Doherty et al., 2022; Niu & Spradling, 2022; Spradling et al., 2022). Particularly, *Drosophila* oocytes provide a perfect system to study, owing to its high accessibility, the amenability to live-cell microscopy, and the ease of genetic manipulation (Peters & Berg, 2016). Over the years, numerous studies in *Drosophila*, demonstrated the importance of MT functions throughout oogenesis and how MT dysregulation leads to defects in oocyte development (Steinhauer & Kalderon, 2006; Lu et al., 2016; Lu et al., 2018; Tillery et al., 2018; Lu et al., 2020).

The *Drosophila* oocyte develops in an egg chamber, where it resides with its 15 germline sister cells, called nurse cells. After the germline stem cell divide, one of the daughter cells undergoes four-round of cell divisions with incomplete cytokinesis, generating an interconnected 16-cell cyst. The oocyte and the nurse cells are connected through cytoplasmic bridges called ring canals, created from incomplete separation during cell divisions. Interestingly, the oocyte is transcriptionally quiescent throughout most of the oogenesis, so its development primarily depends on the nurse cells that transport necessary mRNAs, proteins, and organelles to the oocyte through the ring canals (Cooperstock & Lipshitz, 2001; Bastock & St Johnston, 2008; Hinnant et al., 2020).

In *Drosophila* oogenesis, an oocyte progresses through 14 morphologically distinct developmental stages in an egg chamber. In early oogenesis, centrosomal MTs are nucleated from the oocyte and extend through the ring canals, orienting their plus ends toward nurse cells. The MTs in this array serve as a track for the minus end-directed motor protein, dynein, to transport cargos from nurse cells to the oocyte (Steinhauer & Kalderon, 2006; Tillery et al., 2018; Lu et al., 2023). Then, MTs undergo a dramatic reorganization at stage (ST) 7, at the beginning of mid-oogenesis.

During mid-oogenesis, non-centrosomal MTs in the oocyte nucleate from the anterior cortex of the oocyte, facing away from nurse cells, and form a highly polarized MT gradient within the oocyte (Maro et al., 1990; Theurkauf et al., 1992; Nashchekin et al., 2016; Tillery et al., 2018). This MT array is essential for establishing the polarity of future embryos (Brendza et al., 2000; Cooperstock & Lipshitz, 2001; Steinhauer & Kalderon, 2006; Lu et al., 2020). A recent study revealed that cortical dynein moves MTs in nurse cells against the cortex, generating cytoplasmic flow during late mid-oogenesis (ST9). This cytoplasmic flow carries organelles in bulk from the nurse cells to the oocyte, promoting oocyte growth (Lu et al., 2022).

MT dynamics (polymerization and depolymerization), stability and abundance are regulated by microtubule-associated proteins (MAPs) and post-translational modifications (Baas & Qiang, 2005; Monroy et al., 2020). Notably, it has been shown that about 2% of *Drosophila* genes encode for cytoskeletal proteins and the majority of them remain uncharacterized (Goldstein & Gunawardena, 2000; Karpova et al., 2006). Therefore, we sought to identify new MAPs that regulate MTs during oogenesis.

To find new MAPs, we utilized single-cell RNA (scRNA) data from *Drosophila* ovary reported in a recent study by Jevitt et al. This study reported cell type specific expression of thousands of genes that are expressed in early and mid-oogenesis (Jevitt et al., 2020). After initial analysis of uncharacterized germline-specific genes using Flybase, we found CG18190 as potential new MAP to explore further. The uncharacterized CG18190 encodes a protein of 248 amino acid residues (28.75kDa). FlyAtlas2 (Gillen, A., 2023 Data) reports that CG18190 is highly expressed in the ovary, according to microarray and RNA-Seq data (Chintapalli et al., 2007; Gelbart & Emmert, 2013; Leader et al., 2018). Furthermore, CG18190 was previously included among four *Drosophila* genes with a high degree of sequence similarity to human MT end-binding (EB) protein1 (MAPRE1) (Rogers et al., 2002), suggesting that CG18190 is predicted to be an EB protein.

In this study, we identified a novel microtubule (MT) plus-end tracking protein, by demonstrating its localization at the growing tips of MTs in *Drosophila* S2 cells. Gene expression analysis across tissues reveals that EB-SUN is predominantly expressed in the ovary, whereas EB1 is ubiquitously expressed. The concentrated EB-SUN comets observed in the oocyte during oogenesis reflect high MT polymerization activity, similar to the previously reported enrichment of EB1 comets. EB-SUN knockout (KO) results in reduced MT density at the onset of mid-oogenesis (stage 7) and delays in oocyte growth during late mid-oogenesis (stage 9). While EB-SUN functions redundantly with EB1 during oocyte development, we report that the two EB proteins play distinct roles during early embryogenesis.

## Results

### EB-SUN is a microtubule end-binding protein in Drosophila

To determine whether CG18190 is indeed an EB protein, we utilized Paralog Explorer (Hu et al., 2022) (https://www.flyrnai.org/tools/paralogs/), a recently developed tool for identifying paralogous genes in the fly genome. As expected, EB1 came up, alongside two other genes (CG32371 and CG2955) that encode proteins with more than 50% sequence similarity to CG18190 (Table S1). This result matches the previously identified genes homologous to EB1 (Rogers et al., 2002), suggesting that CG18190 is an EB protein—which we propose to call EB-SUN. Two predicted paralogs, CG32371 and CG2955, are predominantly expressed in testes based on gene expression data from FlyAtlas2 (Chintapalli et al., 2007; Gelbart & Emmert, 2013; Leader et al., 2018), showing low tissue expression correlation. In contrast, EB1 shows the highest tissue expression correlation among those genes (Table S1), suggesting that EB-SUN is likely expressed and functions in the same cells and at the same time as EB1.

Next, we compared amino acid sequences of EB-SUN and EB1 using Clustal Omega (Sievers et al., 2011; Madeira et al., 2024) to identify shared conserved domains (Figure 1A). Residues 17-119 in EB-SUN display the highest degree of conservation within the EB1 family, representing the Calponin Homology (CH) domain essential for MT binding (Juwana et al., 1999; Hayashi & Ikura, 2003). Interestingly, while EB1 has a coiled-coil domain spanning 60 amino acids (residue 213-273), which is crucial for dimerization (Rogers et al., 2002; Slep et al., 2005), EB-SUN features a much shorter coiled-coil region of only 27 amino acids (residue 152-179). Additionally, BLAST searches of EB-SUN sequences against the human genome database revealed that human EB3 (MAPRE3) shows the highest degree of similarity (53%), suggesting that it is likely an orthologue of EB-SUN (Supplemental Figure 1).

**Figure 1.**
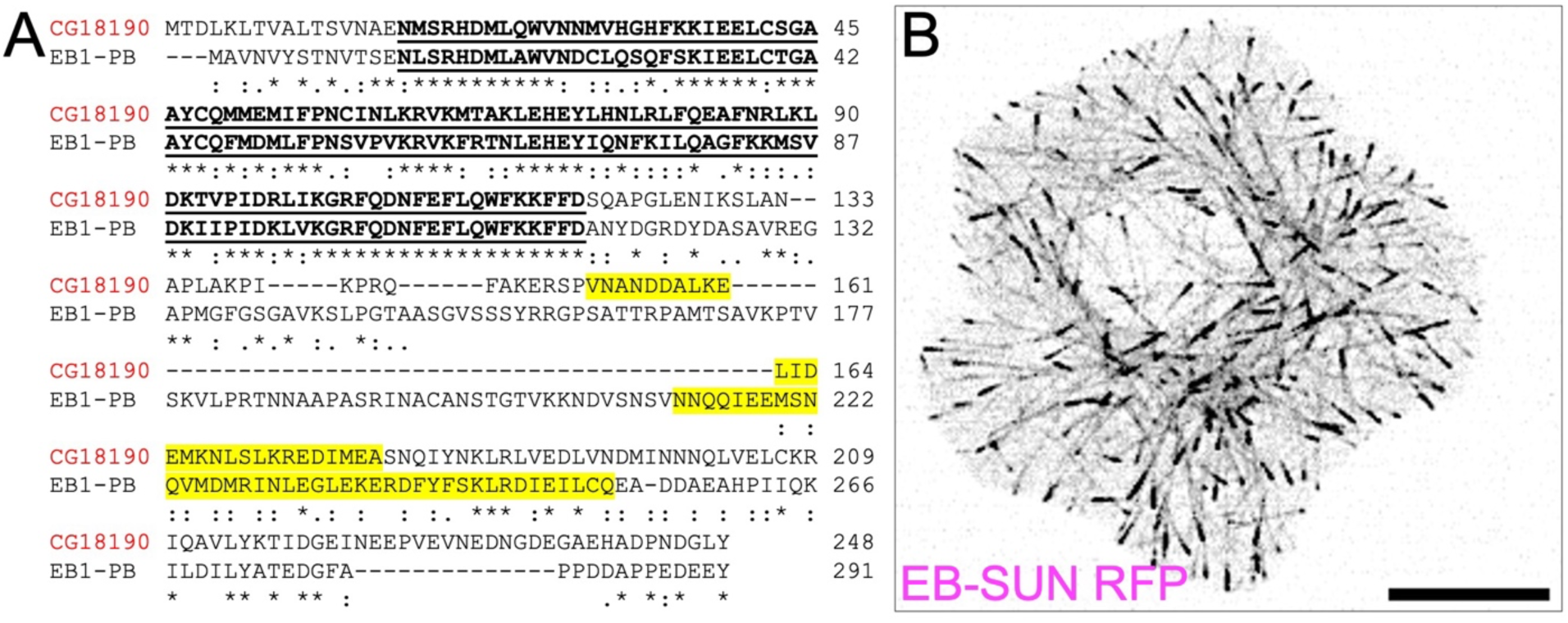
EB-SUN (CG18190) is an EB protein in *Drosophila.* ((A) Protein alignment of EB-SUN with EB1 (isoform PB). The Calponin Homology (CH) domain is indicated by bold, underlined text, and the putative coiled-coil domain is highlighted in yellow. An asterisk (*) indicates residues identical in both sequences; a colon(:) represents conserved substitutions; and a period(.) indicates semi-conserved substitutions (as defined in Clustal Omega). (B) RFP-tagged EB-SUN localizes to the plus ends of microtubules in S2 cells. Scale bar: 10 µm. (See Supplemental Movie S1).

Lastly, we examined the intracellular localization of EB-SUN in S2 cells. Initially, we generated both N- and C-terminally RFP-tagged EB-SUN constructs. While an N-terminal tagged version of EB-SUN was diffusely distributed within cells, C-terminally tagged EB-SUN (EB-SUN-1XTag RFP) formed comets localized to the growing tips of MTs (Figure 1B and Supplemental Movie S1). This confirms that EB-SUN is indeed a new microtubule plus-end tracking protein in *Drosophila*.

### EB-SUN and EB1 colocalize at the growing tips of microtubules

Previous studies of the EB1 crystal structure revealed that EB1 forms a homodimer through a coiled-coil region (Slep et al., 2005). Modeling the structure of EB1 using AlphaFold accurately predicted EB1 homodimerization using fragments from the coiled-coil to the C-terminus (Figure 2A), recapitulating the original crystal structure. Based on the sequence similarity between EB-SUN and EB1, we hypothesized that EB-SUN also forms a homodimer, similar to EB1. Consistent with this hypothesis, AlphaFold predicts that EB-SUN forms a homodimer through interactions within its coiled-coil domain (Figure 2B). Notably, the model reveals a long, unstructured C-terminus in EB-SUN. Unlike the acidic EEY-motif in EB1’s tail, which is crucial for interactions with its binding partners such as CLIP-170 (Komarova et al., 2005; Mishima et al., 2007; Komarova et al., 2009), EB-SUN has a GLY-motif (glycine, leucine, and tyrosine) in its C-terminus. This mostly neutral, unstructured C-terminus region of EB-SUN may contribute to its differential affinities for its binding partners, including MTs and other proteins. Finally, we co-expressed RFP-tagged EB-SUN and EGFP-tagged EB1 constructs in S2 cells (Figure 2D and Supplemental Movie S2), demonstrating colocalization of EB-SUN and EB1 at the growing tips of MTs.

**Figure 2.**
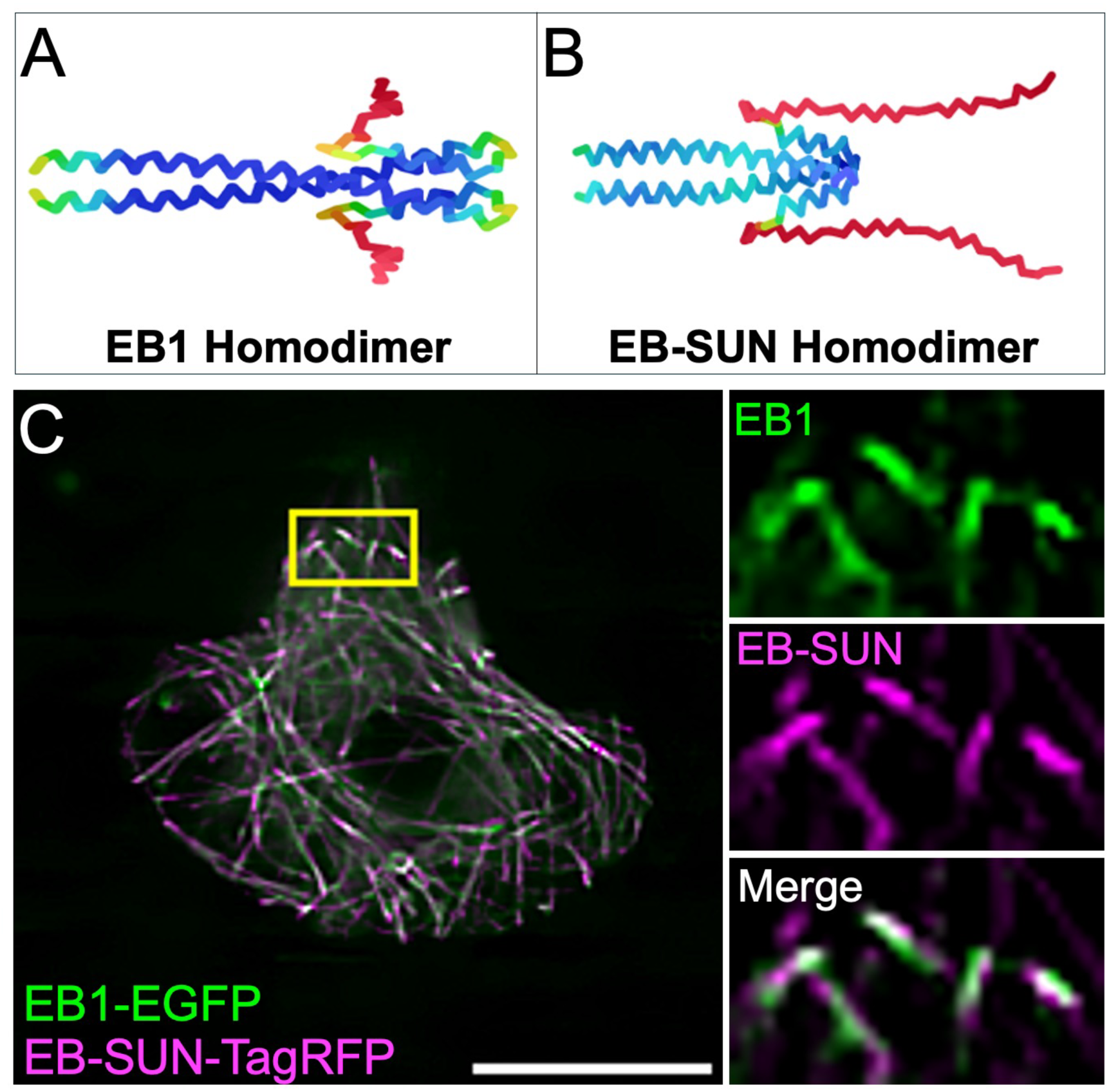
EB-SUN and EB1 localization and structure. (A) AlphaFold accurately predicts the homodimerization of EB1, consistent with previously reported crystal structures. (B) AlphaFold predicts that EB-SUN also forms a homodimer through interactions in the coiled-coil domain, with the N-terminus (The beginning of the coiled-coil domain) oriented to the left and the C-terminus to the right (dark blue for the most confidently predicted regions, via light blue and yellow to orange/red for the regions of very low confidence, as described in AlphaFold). (C) EB-SUN and EB1 colocalize at the growing tips of microtubules in S2 cells. Scale bar: 10 µm. (See Supplemental Movie S2).

### EB-SUN is predominantly expressed in the ovary and enriched in the oocytes during oogenesis

Among the three paralogs of EB-SUN, EB1 shows the highest tissue expression correlation score with EB-SUN (Table S1), suggesting that EB1 and SUN-EB are more likely to be expressed simultaneously. To further investigate their expression patterns across different tissues and developmental stages, we compared EB-SUN and EB1 using Paralog Explorer. This analysis revealed that EB1 is ubiquitously expressed in all tissues (Figure 3A), as shown in human EB1 (Su & Qi, 2001). In contrast, EB-SUN is predominantly expressed in the ovary, which agrees with our initial finding of EB-SUN (CG18190) from scRNA data of the ovary (Jevitt et al., 2020). Additionally, EB-SUN is highly expressed during early embryonic development and in adult females, reflecting its ovary-specific expression (Figure 3B).

**Figure 3.**
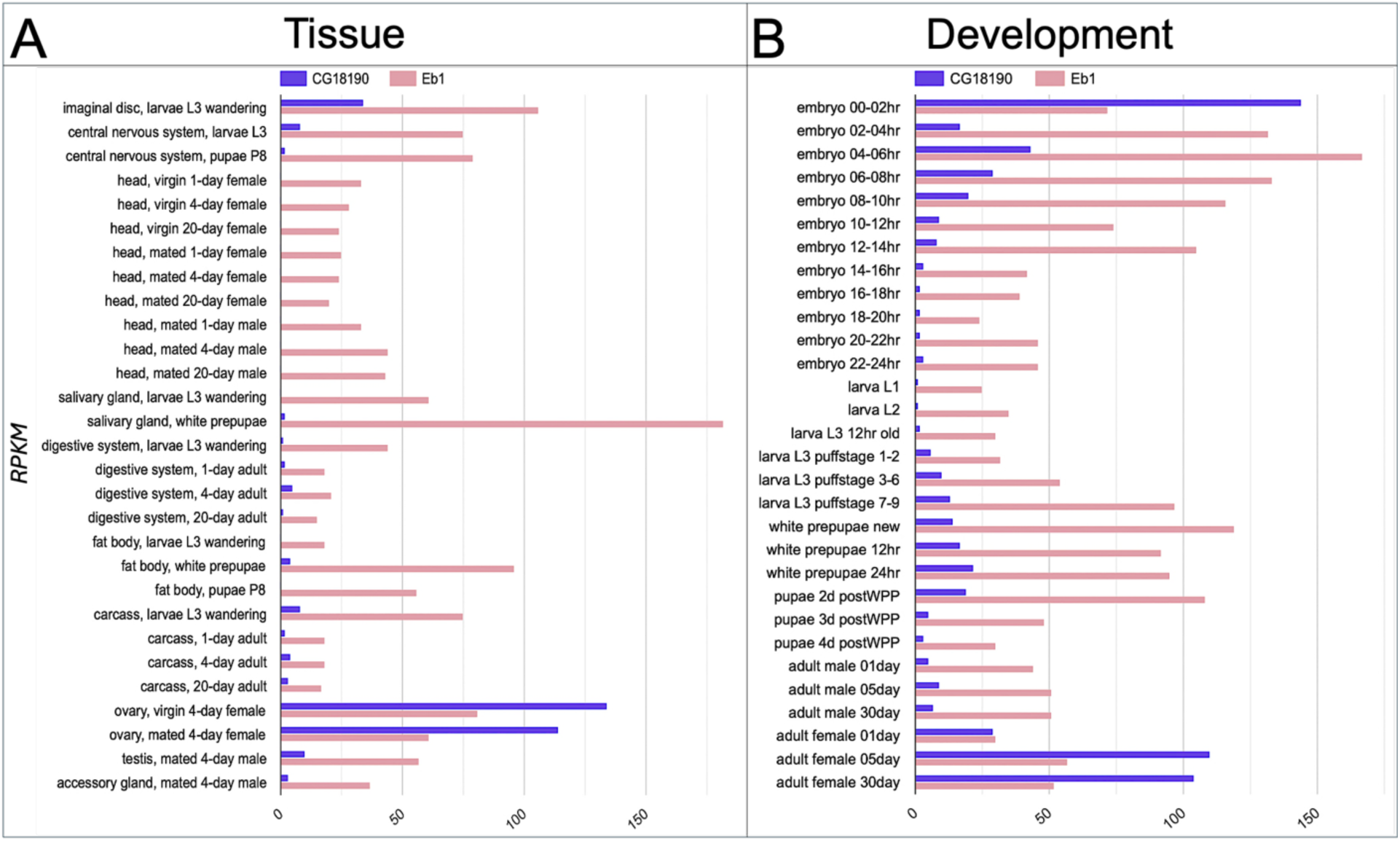
Tissue-specific and developmental expressions of EB-SUN and EB1 in *Drosophila*. (A) Bar graph showing tissue-specific expression levels of EB-SUN and EB1, generated using Paralog Explorer. Unlike EB1, EB-SUN exhibits strong ovary-specific expression (Pearson correlation value: 0.2504). (B) EB-SUN and EB1 appear to be expressed simultaneously during development. EB-SUN is predominantly expressed in early embryos and adult females (Pearson correlation value: 0.1638). Reads Per Kilobase Million (RPKM) is a normalized measure of gene expression used in Paralog Explorer.

A scRNA study reported that EB-SUN is expressed 9-fold more in germline cells than in other cell types within the ovary (Jevitt et al., 2020). Therefore, we examined the localization of EB-SUN *in vivo* during oogenesis. To assess EB-SUN mRNA localization, we employed single-molecule inexpensive fluorescence in situ hybridization (smi-FISH) (Tsanov et al., 2016; Calvo et al., 2021; Lu et al., 2023). We used probes recognizing sfGFP as a negative control and probes targeting Mini spindles (Msps), a MT polymerase concentrated in oocytes during oogenesis (Lu et al., 2023), as a positive control. Our results revealed no specific enrichment of EB-SUN mRNAs (Figure 4A).

**Figure 4.**
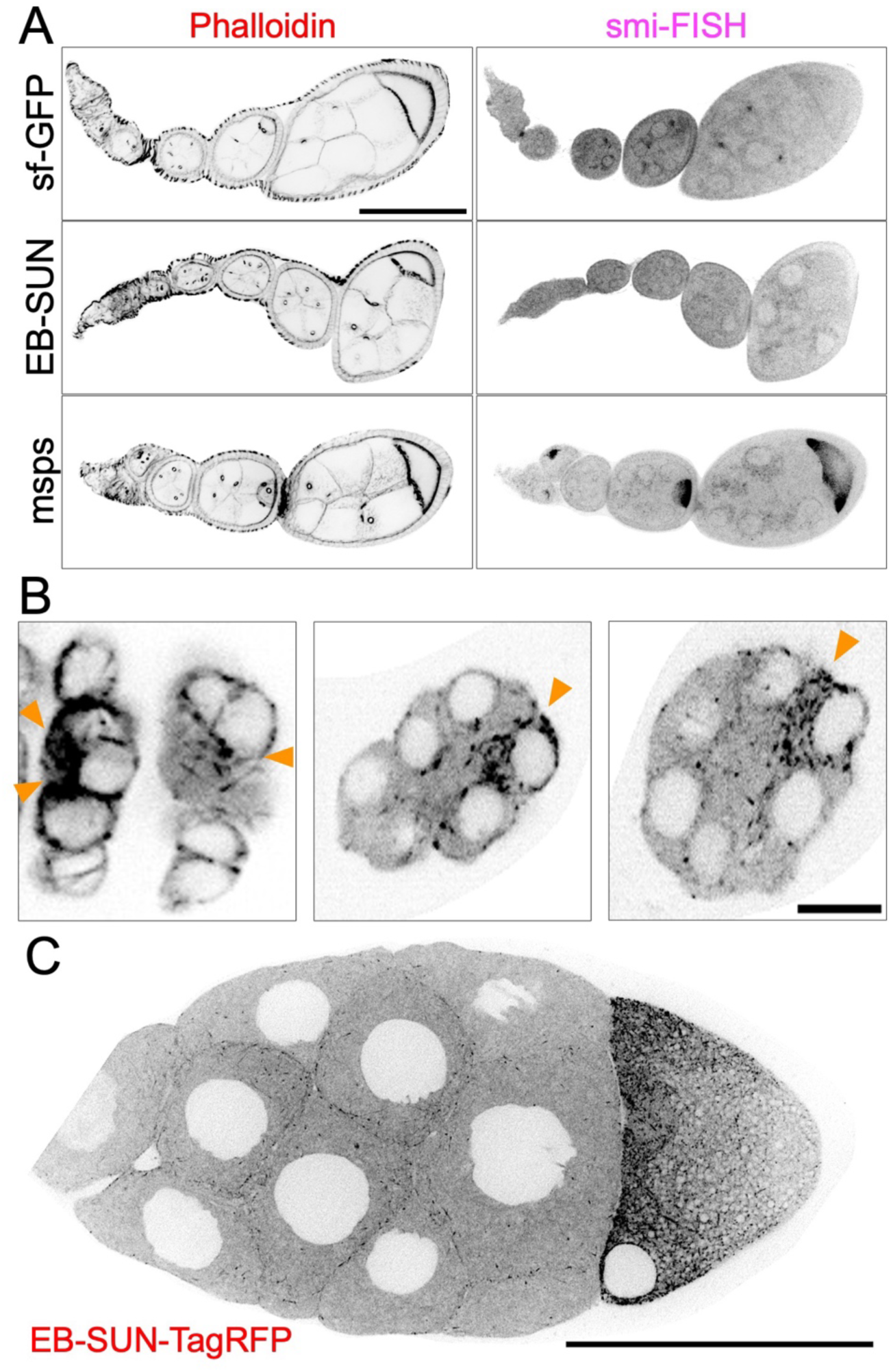
Localization of EB-SUN in *Drosophila* ovary. (A) smi-FISH staining of *sf-GFP* (negative control), *EB-SUN*, and *msps* mRNA (positive control). While *msps* mRNA is concentrated in oocytes as previously shown, there is no enrichment of *EB-SUN* mRNA in oocytes. (B) EB-SUN protein is enriched during oocyte fate determination as early as the two pro-oocyte stage. Scale bar: 10 µm. (See Supplemental Movie S3) (C) During mid-oogenesis, EB-SUN comets are enriched at the anterior cortex of the oocyte, forming a gradient. Scale bar: 100 µm. (See Supplemental Movie S4).

During early and mid-oogenesis, EB1 comets are enriched in oocytes, reflecting a significant increase in MT polymerization activity compared to nurse cells (Lu et al., 2021; Nashchekin et al., 2021; Lu et al., 2023). To observe EB-SUN protein localization, we generated a fly line expressing EB-SUN-1X Tag RFP, under the control of the UAS-Gal4 system (Brand & Perrimon, 1993). We performed live imaging using an early germline driver, *nanos-Gal4^[VP16]^* (Van Doren et al., 1998), which activates EB-SUN-RFP expression in the primordial germ cells. In contrast to EB-SUN mRNA localization, EB-SUN protein comets were enriched in oocytes (Figure 4B and Supplemental Movie S3). Particularly, we observed the enrichment of EB-SUN comets as early as in two pro-oocytes during oocyte fate determination, consistent with previous observations of concentrated EB1 comets in oocytes (Nashchekin et al., 2021; Lu et al., 2023). We also examined EB-SUN localization during mid-oogenesis using a postmitotic germline-specific driver, *maternal α-tubulin-Gal4^[V37]^* (Lu et al., 2021; Lu et al., 2022; Lu et al., 2023). We found that EB-SUN comets are concentrated at the anterior cortex of the oocyte (Figure 4C and Supplemental Movie S4), reflecting the MT gradient critical for establishing polarity (Lu et al., 2020). Collectively, our findings show that EB-SUN comets are enriched in oocytes from early through mid-oogenesis, similar to EB1 localization, suggesting that EB-SUN and EB1 may function redundantly during oocyte development.

### Depletion of EB-SUN reduces MT density and causes a delay in oocyte growth

To investigate the biological function of EB-SUN, we created an EB-SUN knockout (KO) fly line using the CRISPR/Cas9 system. We designed two guide RNAs targeting the beginning of the EB-SUN coding sequence (CDS) and the 3’UTR region. This removed most of the EB-SUN CDS, leaving only two amino acids (see Materials and Methods). The EB-SUN CDS deletion was confirmed by genomic PCR. Additionally, immunoblot analysis of ovary extracts using a polyclonal antibody generated in this study (see Materials and Methods) further confirmed the complete deletion of EB-SUN (Figure 5A and Supplemental Figure S2). Notably, EB1 levels remained unaffected by the EB-SUN KO. Homozygous EB-SUN KO flies are viable and exhibited normal egg-laying activities, suggesting that oocyte development, including germ cell divisions and oocyte specification, was not significantly impacted.

**Figure 5.**
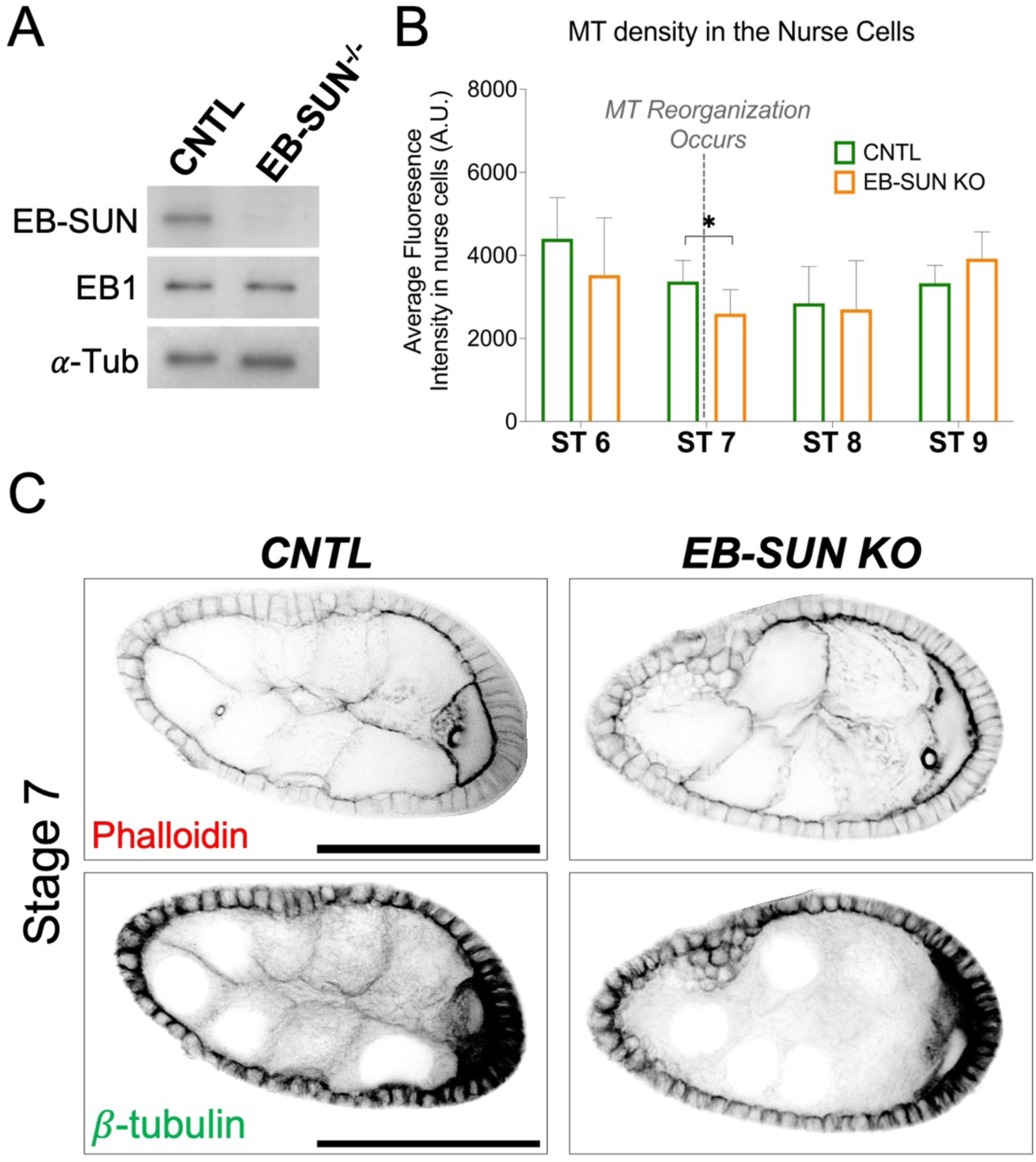
Impact of EB-SUN KO on MTs during oogenesis. (A) Immunoblot of ovary extracts from control and homozygous EB-SUN KO fly lines generated in this study, showing specific depletion of EB-SUN without affecting EB1. *α*-Tubulin antibody was used as a loading control. (B) MT density in the nurse cells was measured. Depletion of EB-SUN causes a decrease in the overall MT density in nurse cells at stage 7 (ST7), where MTs undergo reorganization (p = 0.24 for ST6, p = 0.03 for ST7, p = 0.81 for ST8, p = 0.11 for ST9). Data are represented as the mean with 95% CI. A multiple unpaired t-test with Welch’s correction was performed. For control egg chambers: N = 11 (ST6); N = 11 (ST7); N = 6 (ST8); N = 17 (ST9). For EB-SUN KO egg chambers: N = 12 (ST6); N = 9 (ST7); N = 8 (ST8); N = 12 (ST9). (C) F-actin and β-tubulin staining showing differences in MT density in the nurse cells of control and EB-SUN KO egg chambers at ST7. Scale bar: 50 µm.

While MTs in oocytes are highly dynamic throughout early and mid-oogenesis, as demonstrated by the concentration of EB-SUN and EB1 comets, MTs in nurse cells are stable and exhibit less dynamic instability (Lu et al., 2021; Lu et al., 2022). To investigate the impact of EB-SUN KO on these stable MTs found in nurse cells, we performed immunostaining for β-tubulin and measured MT density in nurse cells. Both control (CNTL) and EB-SUN KO egg chambers were imaged using the same imaging parameters. We observed decreased MT density in the nurse cells at ST 7 (Figure 5, B and C), a critical stage marked by MT reorganization during the transition from early to mid-oogenesis (Tillery et al., 2018). However, overall MT density before and after ST 7 remained largely unaffected by EB-SUN depletion.

Additionally, EB-SUN depletion caused a delay in oocyte growth during late mid-oogenesis, particularly at ST 9 (Figure 6A). Considering ST 9 is a period of significant oocyte growth (Lu et al., 2022), we adopted an analysis method from a recent study (Zhao et al., 2022) to better understand this phenomenon. We subdivided ST 9 into “early” and “late” stages by calculating the ratio of the distance traveled by border cells to the total length of the egg chamber along the anterior-posterior axis (Figure 6B). We then measured the ratio of oocyte area to egg chamber area. Our findings revealed that oocytes from EB-SUN-KO in early ST 9 were smaller than controls (Figure 6D), even though the total egg chamber area remained unchanged (Figure 6E). These data suggest that EB-SUN KO specifically affected oocyte growth. Despite this delay in early ST 9, oocytes eventually caught up to average size by late ST 9 (Figure 6, A and D). While oocyte growth was initially delayed, the overall MT density in nurse cells and the MT gradient in oocytes remained unaffected (Figure 5B and Supplemental Figure S3). Furthermore, homozygous EB-SUN KO females exhibited normal fertility, and their eggs showed normal hatching rates (Figure 6C). These observations suggest that EB-SUN depletion alone does not disrupt major MT dynamics or MT-associated cellular functions, possibly due to redundancy between EB-SUN and EB1.

**Figure 6.**
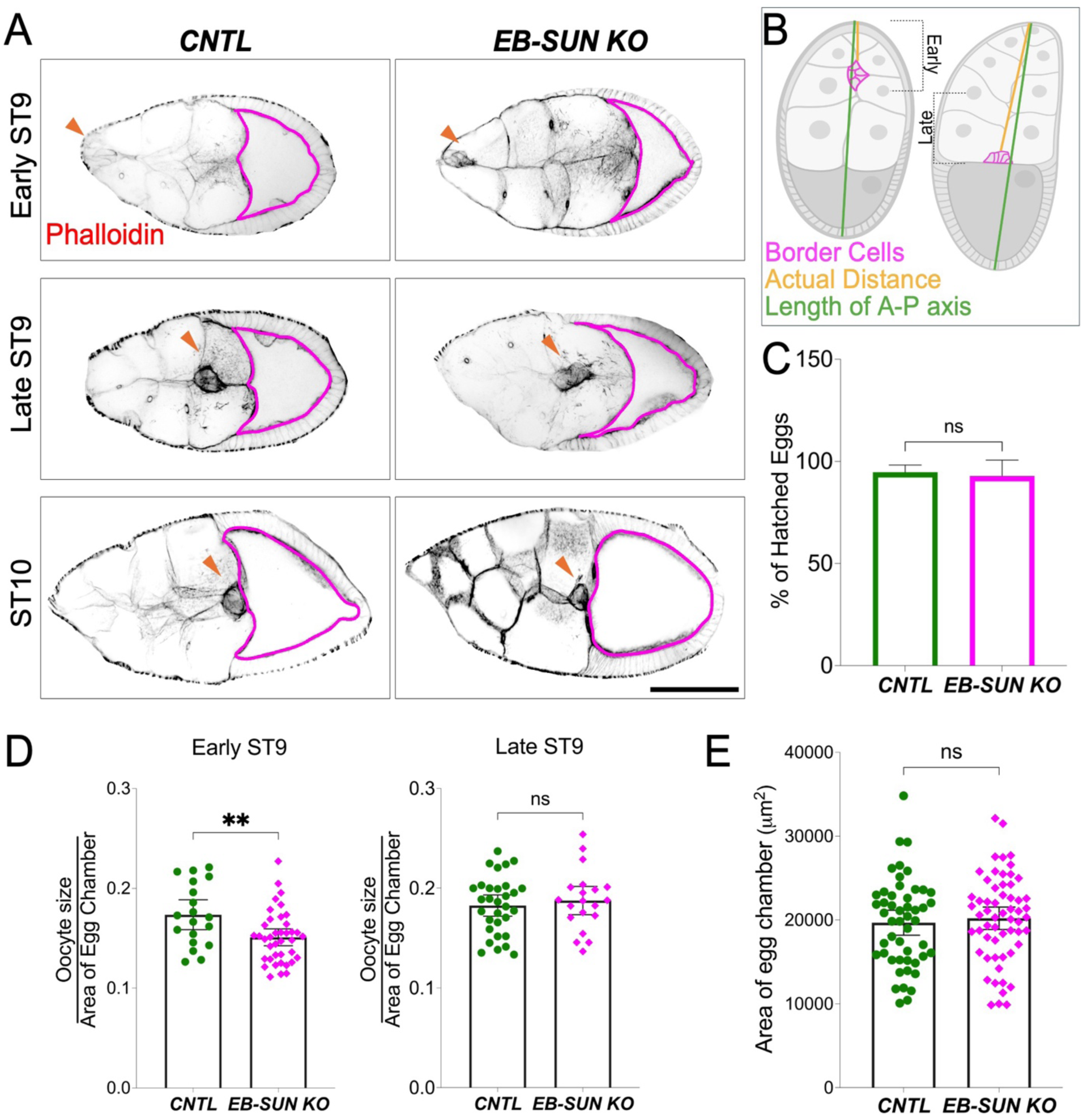
Oocyte growth is delayed in EB-SUN KO during late mid-oogenesis. A) In early ST9, the size of homozygous EB-SUN KO oocytes is smaller than that of controls. However, by late ST9, oocytes in the EB-SUN KO catch up to a size similar to that of controls. Scale bar: 100 µm. (B) Schematic illustrating how early and late stages were defined based on border cell migration. (C) Hatching rate of homozygous EB-SUN KO fly lines. The hatching assay was performed by counting the number of hatched eggs 24 hours after laying. Data are represented as the mean with 95% CI. N = 420 for control; N = 378 for EB-SUN KO from 12 independent collections. Two-tailed unpaired t-tests were performed. (D) The fraction of oocyte size relative to the total area of the egg chamber was measured in early and late ST9. Oocytes in EB-SUN KO are significantly smaller than those in controls at early ST9, but there is no difference at late ST9 (p=0.0053 for early ST9, p=0.561 for late ST9). For controls: N = 19 (early ST9), N = 31 (late ST9); For EB-SUN KO: N = 39 (early ST9), N = 20 (late ST9). (E) Quantification of the egg chamber area at ST9 in both control and EB-SUN KO shows no significant difference. N = 50 for control; N = 59 for EB-SUN KO. Error bars represent 95% CI. Two-tailed unpaired t-tests were performed.

### EB-SUN and EB1 function in a redundant manner during oocyte development

To test whether EB-SUN and EB1 are functionally redundant in oogenesis, we first knockdown (KD) EB1 by expressing EB1-RNAi using both *nanos-Gal4^[VP16]^* and *maternal α-tubulin-Gal4^[V2H]^*, to ensure EB1 depletion in both early germ cells and throughout the oocyte development. We confirmed depletion of EB1 by expressing EB1-RNAi in a GFP-tagged EB1 background, showing EB1 comets are abolished in germline cells (Supplemental Figure 4 and Supplemental Movie S5). As we observed in homozygous EB-SUN KO females, there were no significant phenotypes related to cell divisions or oocyte specification, and the overall oogenesis appeared normal (Supplemental Figure 5). These findings indicate their functional redundancy during oogenesis, including in oocyte specification and development.

Next, we investigated the impact of double EB-SUN KO/EB1 KD on oocyte development. To bypass the requirement for one of EB proteins in early cell divisions and oocyte specification, we used a postmitotic germline-specific driver, *maternal α-tubulin-Gal4^[V37]^* (Lu et al., 2021; Lu et al., 2022; Lu et al., 2023). This approach allowed us to express EB1-RNAi in homozygous EB-SUN KO flies after completing early cell division and oocyte determination. We first examined overall MTs in nurse cells and found that EB1 KD alone caused a decrease in MT density at ST 7 (Figure 7, A and C), similar to what we observed in EB-SUN KO (Figure 5, B and C). Moreover, the double depletion of both EB proteins led to a significantly greater reduction in MT density at stage 7 (Figure 7, A and C). Surprisingly, there are still MTs remaining in nurse cells upon double depletion, which can be explained by previous studies showing MTs in nurse cells are extremely stable (Lu et al., 2022; Lu et al., 2023).

**Figure 7.**
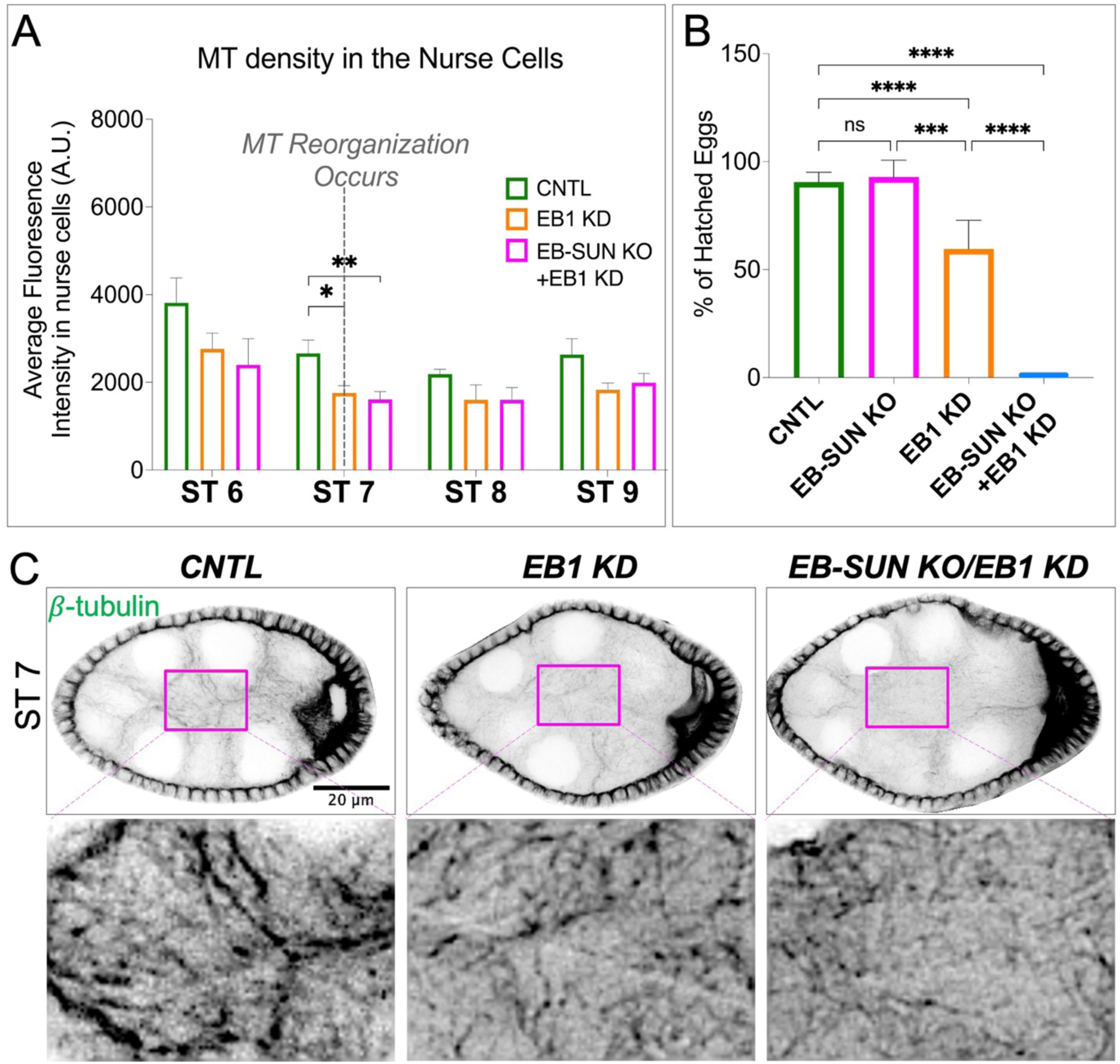
Impact of EB-SUN KO/EB1 KD on MTs during oogenesis and embryogenesis. (A) MT density in the nurse cells was measured. Depletion of both EB-SUN and EB1 causes a significant decrease in the overall density of MTs in the nurse cells at ST 7, where MTs undergo reorganization. Data are represented as mean with 95% CI. One-way ANOVA with Tukey’s multiple comparisons test was performed for each stage (p=0.027 for EB1 KD; p=0.0056 for EB-SUN KO/EB1 KD at ST 7). For control egg chambers: N=4 ST6; N=7 ST7; N=7 ST8; N=6 ST9. For EB1 KD egg chambers: N=6 ST6; N=8 ST7; N=3 ST8; N=7 ST9. For EB-SUN KO/EB1 KD egg chambers: N=6 ST6; N=11 ST7; N=9 ST8; N=10 ST9.(B) Quantification of the number of eggs that hatched 24 hours after laying. Data are represented as mean with 95% CI. N=522 for control; N=378 for EB1 KD; N=91 for EB-SUN KO/EB1 KD from 14 independent collections. Multiple unpaired t test with Welsh correction was performed. (C) β-tubulin staining showing differences in MT density in the nurse cells of control, EB1 KD alone, and double EB-SUN KO/EB1 KD egg chambers. (The contrast of insets showing the enlarged view of MT density was increased).

Lastly, we conducted a hatching assay by counting the eggs hatched 24 hours after laying. EB1 KD alone significantly decreased the hatching rate (Figure 7B). This observation is consistent with a previous study showing that EB1 inhibition causes defective spindle elongation and chromosomal segregation (Rogers et al., 2002). Notably, the double depletion of both EBs resulted in a 0% of hatching rate, demonstrating the redundant functions of EB-SUN and EB1 in early embryogenesis. Intriguingly, EB1 KD alone affected embryogenesis, whereas homozygous EB-SUN KO flies exhibited a normal hatching rate (Figure 6C and 7B). These findings suggest that although, both EBs are expressed in early embryos (Figure 3), EB1 and EB-SUN may be differentially regulated or play distinct roles during early embryogenesis.

## Discussion

Microtubules are essential during *Drosophila* oogenesis, undergoing stereotypical changes in stability, abundance, and biological function throughout the process. Here, we identified an uncharacterized gene, CG18190, initially found in scRNA data from the *Drosophila* ovary. Paralog analysis of CG18190 against both *Drosophila* and human genomes revealed that *Drosophila* EB1 and human EB1 (MAPRE3) are paralogs of EB-SUN, with a high degree of similarity (50%). Additionally, we observed its localization at the growing MT tips in S2 cells. Both paralog analysis and intracellular localization confirm that CG18190 is a new MT plus-end tracking protein in *Drosophila,* and we propose to name it EB-SUN.

### Structural Features of EB-SUN

Comparing the amino acid sequences of EB-SUN and EB1 revealed that both proteins contain the CH and EB1 homology domains, which are highly conserved within the EB family (Akhmanova & Steinmetz, 2008). Additionally, compared to EB1’s coiled-coil domain, which is required for dimerization, EB-SUN possesses a much shorter coiled-coil domain. Mammalian cells have three EB proteins and EB1(MAPRE1) and EB3 (MAPRE3) have been shown to heterodimerize and rapidly exchange with one another (De Groot et al., 2010). Based on the sequence similarity between EB-SUN and EB1 and evolutionarily conservation, we assume that EB-SUN forms a heterodimer with EB1. However, a limitation of this structural analysis is that we used only fragments from the coiled-coil domain to the C-terminus rather than full sequences because the flexible domain between the coiled-coil and CH domains made the models difficult to interpret. Notably, the full-length protein prediction for EB-SUN showed that its N-terminal domain is quite different from EB1 and has a long, unstructured C-terminus. We speculate that these regions may confer different binding affinities or enable unique protein-protein interactions compared to EB1.

### Expression Patterns of EB-SUN and EB1

In mammalian cells, numerous studies demonstrated that EB proteins are differentially regulated and have different cellular functions. For example, when all three EB proteins are depleted, only EB1 and EB3, but not EB2, can promote persistent microtubule growth and restore CLIP-170 localization to the growing MT tips (Komarova et al., 2005; Komarova et al., 2009). Additionally, EB2 is highly expressed during epithelial polarization (Goldspink et al., 2013) and is key in regulating focal adhesions in migrating cells (Yue et al., 2014). Comparisons of EB-SUN and EB1 expression across fly tissues and developmental stages using Paralog Explorer revealed that EB1 is ubiquitously expressed across all tissues and developmental stages. This finding is consistent with observations in human EB1 (MAPRE1) (Su & Qi, 2001). In contrast, EB-SUN exhibits ovary-specific expression and is highly expressed during early embryonic stages. This distinct expression pattern raises intriguing questions about the role of EB-SUN in the ovary. Could it have distinct functions or unique binding partners in the ovary, separate from those of EB1? Pull-down assays with ovary extracts using recombinant EB proteins or proximity labeling experiments can be conducted to identify unique binding partners for EB1 and EB-SUN. This could provide insight into distinct molecular pathways in which they function, as shown in mammalian EBs (Komarova et al., 2005; Komarova et al., 2009; Goldspink et al., 2013; Yue et al., 2014).

### Functional Analysis of EB-SUN in Oogenesis

To study the function of EB-SUN during oocyte development, we deleted the CDS using CRISPR/Cas9 to generate an EB-SUN KO allele. EB-SUN depletion reduced MT density in nurse cells at ST 7, the onset of mid-oogenesis. Immunostaining revealed that most MT reduction in EB-SUN KO nurse cells is due to the loss of cortical-enriched MTs, which are prominent in control but absent in the EB-SUN KO line (Figure 5C). This is consistent with the observation that EB-SUN comets in nurse cells are predominantly found near the cell cortex (Figure 4C and Supplemental Movie S4). Despite the loss of this cortical MT, the larger pool of MTs within the cytosol of nurse cells remained largely unaffected, suggesting the presence of two distinct MT pools: 1) highly dynamic MTs near the cortex and 2) MTs at steady-state (i.e., stable MTs) within the cytosol. We also observed that EB-SUN KO caused a delay in oocyte growth in the early ST 9. However, the oocytes reached normal size by late ST 9. We speculate that the cytoplasmic flow at stage 9—a transport mechanism in which organelles move in bulk from the nurse cells to the oocyte (Lu et al., 2022), enables oocytes to compensate for slower initial rates of growth.

### Functional Redundancy Between EB-SUN and EB1

To investigate the functional redundancy of EB-SUN and EB1 during oocyte development, we first depleted EB1 alone. Similar to the observations in the EB-SUN KO, EB1 KD led to a reduction in MT density at ST 7 (Figure 7, A and C), primarily affecting cortical MTs in nurse cells. However, no defects were observed in overall oogenesis or egg-laying activity. The same localization of both EB proteins (i.e., enrichment in oocyte and cortical localization in nurse cells) and the similar effects of their depletion on MTs in nurse cells suggest functional redundancy between EB-SUN and EB1 during oogenesis.

Our results from the double depletion of EB-SUN and EB1 showed a further reduction in MT density at ST7, compared to the depletion of either EB protein alone (Figure 7C). Surprisingly, we did not observe any significant MT-associated defects, such as small oocytes or the absence of a MT gradient for polarity establishment (Theurkauf et al., 1992; Steinhauer & Kalderon, 2006; Lu et al., 2020; Lu et al., 2021, 2023). Furthermore, overall oogenesis appeared normal, and the eggs were fully developed. Oocyte development relies on MT-dependent transport of all necessary organelles from nurse cells (Lu et al., 2021; Lu et al., 2022). Despite the reduction in MTs in nurse cells upon double EB-SUN KO and EB1 KD, our findings suggest that the depletion of both EB proteins did not severely impact MT-dependent transport.

A previous study demonstrated that high MT polymerization activity in oocytes throughout early and mid-oogenesis is necessary for oocyte development (Lu et al., 2023). This MT polymerization activity in oocytes correlates with the presence of EB-SUN or EB1 comets we observed (Lu et al., 2021). However, based on normal MT arrays we observed throughout oogenesis via immunostaining of β-tubulin, we speculate that both EB proteins may not be essential for MT polymerization.

### Distinct Roles in Early Embryogenesis

Intriguingly, we found that EB1 KD alone significantly reduced the hatching rate, in contrast to the normal hatching rate observed in EB-SUN KO (Figure 7B). This finding suggests differential regulation or distinct functions between EB-SUN and EB1 during early embryogenesis. *Drosophila* embryos undergo 13 rounds of syncytial divisions (Deneke et al., 2019). Previously, EB1 inhibition in syncytial embryos disrupted the elongation of the mitotic spindle, preventing proper chromosome segregation (Rogers et al., 2002). This disruption likely explains the decreased hatching rate observed in EB1 KD alone (Figure 7B). Apart from the functional redundancy between the two EB proteins, as demonstrated by the severely reduced hatching rate (0%) in double EB-SUNKO/EB1 KD, further work testing whether overexpression of EB-SUN can compensate for the loss of EB1 function during embryogenesis could provide valuable insights.

### Conclusion and Future Directions

In conclusion, our studies reveal that EB-SUN KO or EB1 KD alone does not cause major MT-associated phenotypes during oogenesis. Additionally, we demonstrated that EB-SUN and EB1 are functionally redundant in oocyte development, but our findings suggest that these two EB proteins have distinct roles in early embryogenesis. Notably, the ubiquitous expression of EB1 and the tissue-specific expression of EB-SUN in *Drosophila* resemble the expression patterns of EB proteins in humans. While this study focused on EB-SUN in the ovary, further research exploring the two other predicted paralogs of EB-SUN (CG32371 and CG2955), which exhibit testes-specific expression, would be interesting. Our study of EB-SUN, along with the generation of a fly line expressing the EB-SUN-1X Tag RFP transgene, provides an excellent tool for further research into the distinct properties of these EB paralogs, leveraging the advantages of the *Drosophila* model system.

## Acknowledgments

We thank all the members of the Gelfand laboratory for their support, discussions, and suggestions. Special thanks to our summer research intern, Anisha Verma. We also thank the Northwestern Center for Advanced Microscopy & Nikon Imaging Center for imaging assistance, and the Northwestern Sanger Sequencing Facility for sequencing services. Research reported in this study was supported by the National Institute of General Medical Sciences grant R35GM131752 to V.I.G. We also acknowledge the Bloomington *Drosophila* Stock Center (supported by NIH grant P40OD018537) for providing fly stocks.

## Materials and Methods

### Drosophila strains

Fly stocks and crosses were maintained on standard cornmeal food (Nutri-Fly Bloomington Formulation, Genesee, Cat #: 66–121) supplemented with dry active yeast at room temperature (∼24–25°C). *The UASp-EB-SUN (CG18190)-1X Tag RFP fly line was generated in this study using PhiC31-mediated integration (BestGene Inc.).* The following fly stocks were used in this study: *mat αtub-Gal4^[V37]^* (III, Bloomington *Drosophila* Stock Center #7063); *nos-Gal4^[VP16]^* (III) (Van Doren et al., 1998; Lu et al., 2012); *ubi-EB1-GFP* (III) (from Dr. Steve Rogers, the University of North Carolina at Chapel Hill) (del Castillo et al., 2015; Lu et al., 2015, 2021); *UAS-EGFP-RNAi* (BDSC#: 41551); *UAS-EB1 RNAi* (GL00559 attP2, III, BDSC# 36599, targeting CDS 5’-TCGGTCAACAATCAACAAATA-3’); *UAS-EB1 RNAi* (HMS01568 attP40, II, BDSC# 36680, targeting EB1’s 3’ UTR region 5’-ATGGTTTACATGGTTGATTTA-3’).

### EB-SUN CRISPR KO Fly Line

The EB-SUN KO line was generated by WellGenetics, Inc. using CRISPR/Cas9-mediated mutagenesis, following modified methods from (Kondo & Ueda, 2013). Briefly, the upstream sequence CATATTCTCCGCATTAACGC[TGG], targeting the beginning of the CDS, and the downstream sequence TGATGAAGGCGCCGAACACG[CGG], targeting the 3’ UTR region, were separately cloned into a U6 promoter plasmid. The RMCE-3xP3-RFP cassette, which contains forward attP, reverse attP, a floxed 3xP3-RFP, and two homology arms, was cloned into the pUC57-Kan vector as a donor template for repair. Microinjection of all DNA plasmids, including gRNAs, hs-Cas9, and the RMCE-3xP3-RFP cassette donor DNA, was performed into embryos of the control strain [LWG228] w[1118]. F1 flies carrying the 3xP3-RFP selection marker were further validated by genomic PCR and sequencing. CRISPR generated an 805-bp deletion, replacing most of the CDS with the RMCE-3xP3-RFP cassette in the EB-SUN allele.

### EB-SUN Polyclonal Antibody

A polyclonal rabbit antibody against CG18190 was generated by FabGennix International Inc. A peptide corresponding to residues 202-230 of EB-SUN(QLVELCKRIQAVLYKTIDGEINEEPVEVN) was synthesized, and the antibodies were affinity-purified.

### Plasmid constructs

pMT.EB1-GFP was provided by Dr. Steve Rogers at the University of North Carolina at Chapel Hill. The pMT.EB-SUN-RFP plasmid was generated in this study. The EB-SUN gene (FlyBase ID#: FBgn0034403) was synthesized by Twist Bioscience. EB-SUN with 2xGGSG linkers and RFP was cloned into the pMT/V5-His vector at the NotI and XbaI sites. The UASp-EB-SUN-1xTag RFP plasmid was also generated in this study, with the EB-SUN-1xTagRFP fragment inserted into the UASp vector (Lu et al., 2022) at NotI and XbaI sites.

### Drosophila S2 cell

*Drosophila* S2R+ cells (DGRC Stock Number: 150) were maintained in Insect-Xpress medium (Lonza, Cat#: BELN12-730Q). S2 cells were transfected with 0.5 µg of DNA in a 12-well plate using the Effectene transfection kit (Qiagen, Cat. #/ID: 301425) and plated 48 hours after transfection on acid-washed No. 1.5 coverslips (Corning). The coverslips had been treated with a solution of 0.5 mg/ml concanavalin A (Sigma-Aldrich) in water and allowed to air dry (Rogers et al., 2002; Rogers & Rogers, 2008). Cells were imaged using a Nikon W1 spinning disk confocal microscope (Yokogawa CSU with 50 µm pinhole size), equipped with a Photometrics Prime 95B sCMOS camera and a 100× 1.45 N.A. oil immersion lens, controlled by Nikon Elements software. EB-SUN or EB1 comets were imaged every 2 seconds for 2 minutes.

### Single-Molecule Inexpensive Fluorescence in situ Hybridization (smiFISH)

Twenty-base-long DNA probes complementary to the mRNA of EB-SUN, with a 3′ FLAP-X complementary probe (5′-CCTCCTAAGTTTCGAGCTGGACTCAGTG-3′), were designed using LGC Biosearch Technologies’ Stellaris RNA FISH Probe Designer (masking level five, minimal spacing of two bases). Probes specific to *sfGFP* were used as a control, and probes targeting *msps* 3′ UTR mRNA were used as a positive control to ensure the functionality of smiFISH (Lu et al., 2023) (probe sequences are listed in Supplemental Table S2). The probes were diluted to 100 μM in nuclease-free H2O. Probes were mixed in equal molar ratios to a final concentration of 100 μM and stored at −20°C. The fluorescently labeled Flap-X probe was ordered from IDT (100 nmol synthesis scale, HPLC purified) and diluted in nuclease-free H2O to a concentration of 100 μM. The mRNA-FLAP-X complementary probes and fluorescent Flap-X probes were annealed by mixing 2 μL of the mixed mRNA-FLAP-X complementary probe (100 μM mixed concentration), 2.5 μL of Cy5-FlapX probe (100 μM), 5 μL of New England Biolabs Buffer 3, and 40.5 μL of nuclease-free H2O. The mixture was incubated at 85°C for 3 minutes, 65°C for 3 minutes, and 25°C for 5 minutes in a PCR machine, and then stored at −20°C. Then, smi-FISH was performed following the protocol as previously described (Lu et al., 2023).

### Live Imaging of Drosophila Egg Chambers

oung mated female adults were fed dry active yeast for 16–18 hours and then dissected in Halocarbon oil 700 (Sigma-Aldrich, Cat# H8898), as previously described (Lu et al., 2020; Lu et al., 2021; Lu et al., 2022). Fluorescent samples were imaged using a Nikon W1 spinning disk confocal microscope (Yokogawa CSU with a 50 µm pinhole size), equipped with either a Photometrics Prime 95B sCMOS camera or a Hamamatsu ORCA-Fusion Digital CMOS camera, and a 40× 1.30 N.A. oil immersion lens or a 40× 1.25 N.A. silicone oil lens, controlled by Nikon Elements software.

### Microtubule Staining of Drosophila Egg Chambers

We used a well-established staining protocol from our lab (Lu et al., 2020; Lu et al., 2021; Lu et al., 2022). Ovaries were dissected in 1× Brinkley Renaturing Buffer 80 [BRB80, 80 mM piperazine-N,N’-bis(2-ethanesulfonic acid) (PIPES), 1 mM MgCl₂, 1 mM EGTA, pH 6.8] and fixed in 8% EM-grade formaldehyde (Electron Microscopy Sciences 16% Paraformaldehyde Aqueous Solution, Fisher Scientific, Cat# 50-980-487) + 1× BRB80 + 0.1% Triton X-100 for 20 minutes on a rotator at room temperature. Samples were briefly washed with 1× PBTB (1× PBS + 0.1% Triton X-100 + 0.2% BSA) five times for 10 minutes each, then blocked in 5% normal goat serum-containing 1× PBTB for 1 hour at room temperature. Samples were stained with CoraLitePlus 488-conjugated or CoraLite 594-conjugated β-tubulin monoclonal antibody (ProteinTech, Cat# CL488-66240, Clone No. 1D4A4; Cat# CL594-66240, Clone No. 1D4A4) at 1:100 dilution at 4°C overnight. Samples were also stained with rhodamine-conjugated or Alexa Fluor 633-conjugated phalloidin (0.2 µg/mL) and DAPI (1 µg/mL) for 1 hour before mounting. Imaging was performed using a Nikon W1 spinning disk confocal microscope (Yokogawa CSU with a 50 µm pinhole size), equipped with a Hamamatsu ORCA-Fusion Digital CMOS camera and a 100× 1.35 N.A. silicone oil immersion lens, controlled by Nikon Elements software. Z-stack images were acquired with a step size of 0.5 µm.

### Measurement of Nurse Cell and Oocyte Size

Using Phalloidin (F-actin) and DAPI channels in Z-stack images of the ovarioles, the nurse cell and oocyte areas (at the largest cross-section) were measured by manually tracing the egg chamber boundaries with the polygon selection tool in FIJI. For stage 9 egg chambers, we adopted the analysis method from a recent study (Zhao et al., 2022). We subdivided stage 9 into “early” and “late” stages by calculating the ratio of the distance migrated by the border cells to the total length of the anterior-posterior axis of the egg chamber. The ratio of the oocyte area to the total egg chamber area was then measured and plotted.

### Quantification of Microtubule Staining in Drosophila Egg Chambers

A sum Z-projection of a 2.5 µm z-stack image (six z-slices in total) from stage 6 to stage 9 egg chambers was created using FIJI (Image > Stacks > Z projection > Sum slices). To ensure unbiased data analysis of microtubule density in nurse cells, a circle with an area of 300×300 pixels was used to measure different regions within nurse cells three times, and the measurements were averaged. Fluorescence intensity in the nucleus was also measured as background and subtracted from the mean MT intensity in the nurse cells. Egg chamber stages were defined based on the characterization previously described (Jia et al., 2016; Lu et al., 2021, 2023).

### Statistical Analysis

The figures show either percentages of phenotypes, mean values, or all data points, as indicated in the figure legends. Error bars represent the 95% confidence interval (CI), and *N* refers to the number of samples examined in each assay. Statistical analysis was performed using GraphPad Prism 8.0.2. *P* values were calculated using unpaired t-tests with Welch’s correction. The levels of statistical significance were assigned as follows: *P* ≥ 0.05, not significant (n.s.); 0.01 ≤ *P*< 0.05, significant (); 0.001 ≤ P < 0.01, very significant (); 0.0001 ≤ P < 0.001, extremely significant (); P < 0.0001, highly significant (****).

---

**Table S1.** Genes that encode proteins with a high degree of sequence similarity (>50%) to EB-SUN (CG18190) are identified via Paralog Explorer.

| **Paralogs of EB-SUN (CG18190)** |  |  |  |  |  |
| --- | --- | --- | --- | --- | --- |
|  | Microarray<br>& RNA-Seq<br>From FlyAtlas2 | Alignment<br>Length | Similarity | Common GO Slim | Tissue<br>Expression<br>Correlation |
| **EB1** |  | 284 | 51% | GO:0043234<br>(proteincomplex);<br>GO:0005622<br>(intracellular);<br>GO:0005856<br>(cytoskeleton);<br>GO:0043226<br>(organelle);<br>GO:0008092<br>(cytoskeletal protein<br>binding);<br>GO:0005623(cell) | 0.25037 |
| **CG32371** | Very High<br>expression<br>in Testes | 180 | 59% | GO:0008092<br>(cytoskeletal protein<br>binding) | 0.00695 |
| **CG2955** | Very High<br>expression<br>in Testes | 137 | 56% | GO:0008092<br>(cytoskeletal protein<br>binding) | 0.05561 |

**Table S2.**
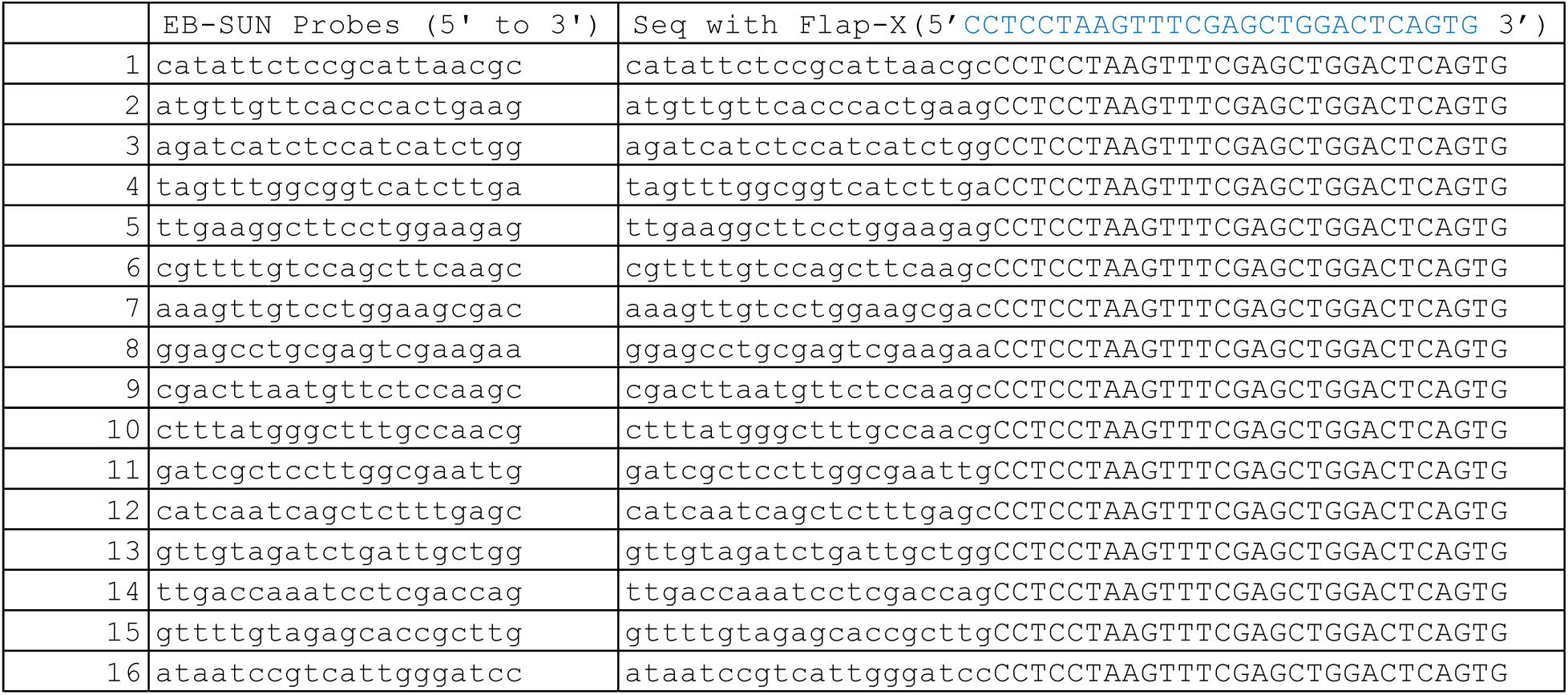
Probes specific to EB-SUN CDS for smi-FISH.

**Figure S1.**
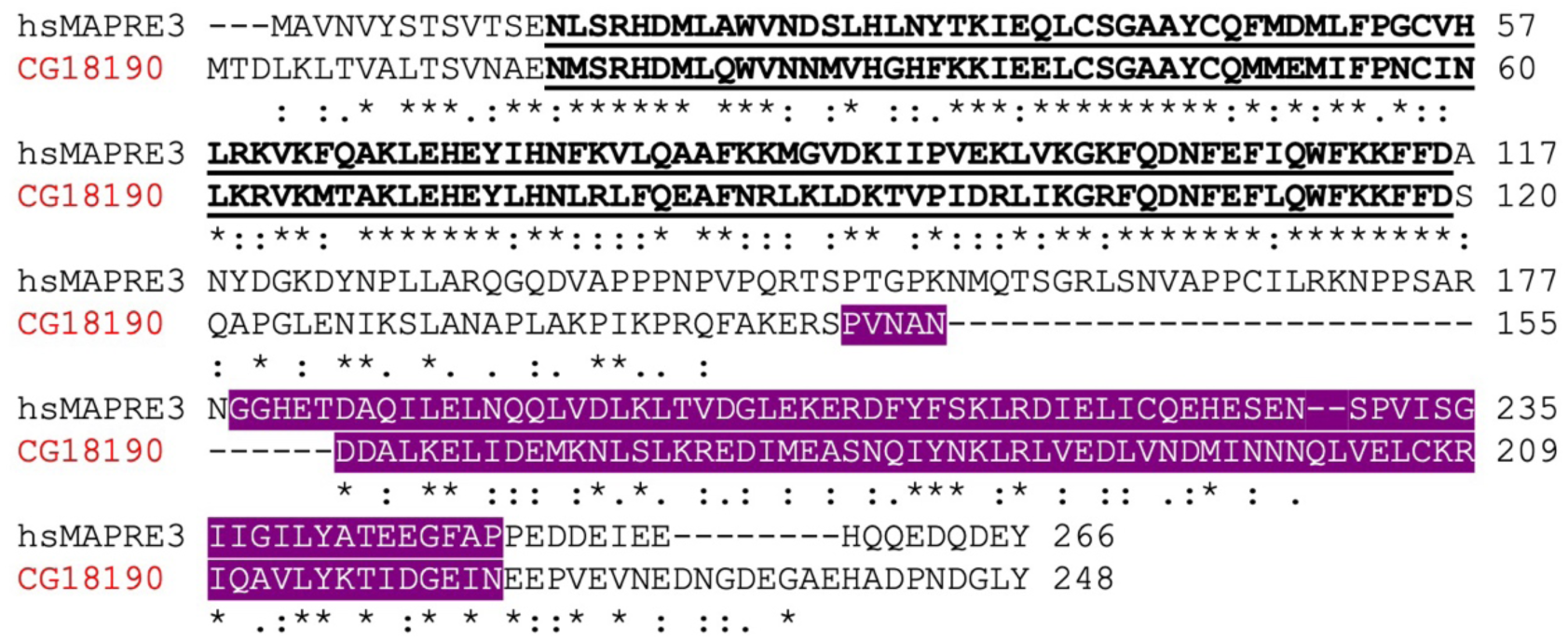
Protein alignment of EB-SUN with human microtubule-associated protein family member 3 (MAPRE3). Underlined text in bold indicates the CH domain (14-116 in MAPRE3 and 17-119 in EB-SUN), and white texts highlighted in purple represent the EB homology domain (194-264 in MAPRE3 and 151-223 in EB-SUN). Asterisk (*), residues that are identical in two sequences in the alignment: colon (:), conserved substitutions. period (.), semi-conserved substitutions (as defined in Clustal Omega).

**Figure S2.**
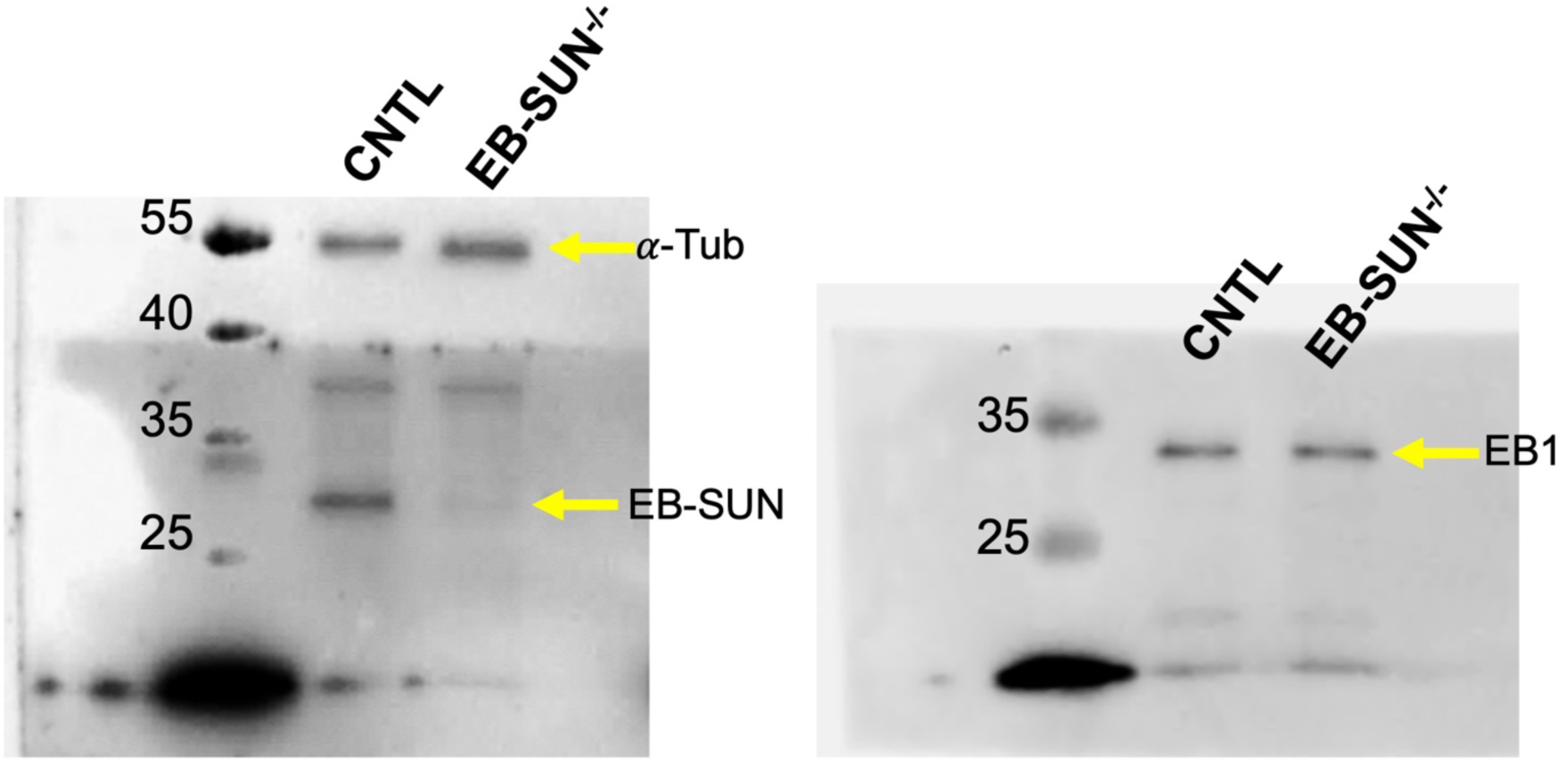
Immunoblot with ovary extracts of CNTL (W^1118^) and homozygous EB-SUN KO. The same PVDF membrane was stripped then probed for EB1.

**Figure S3.**
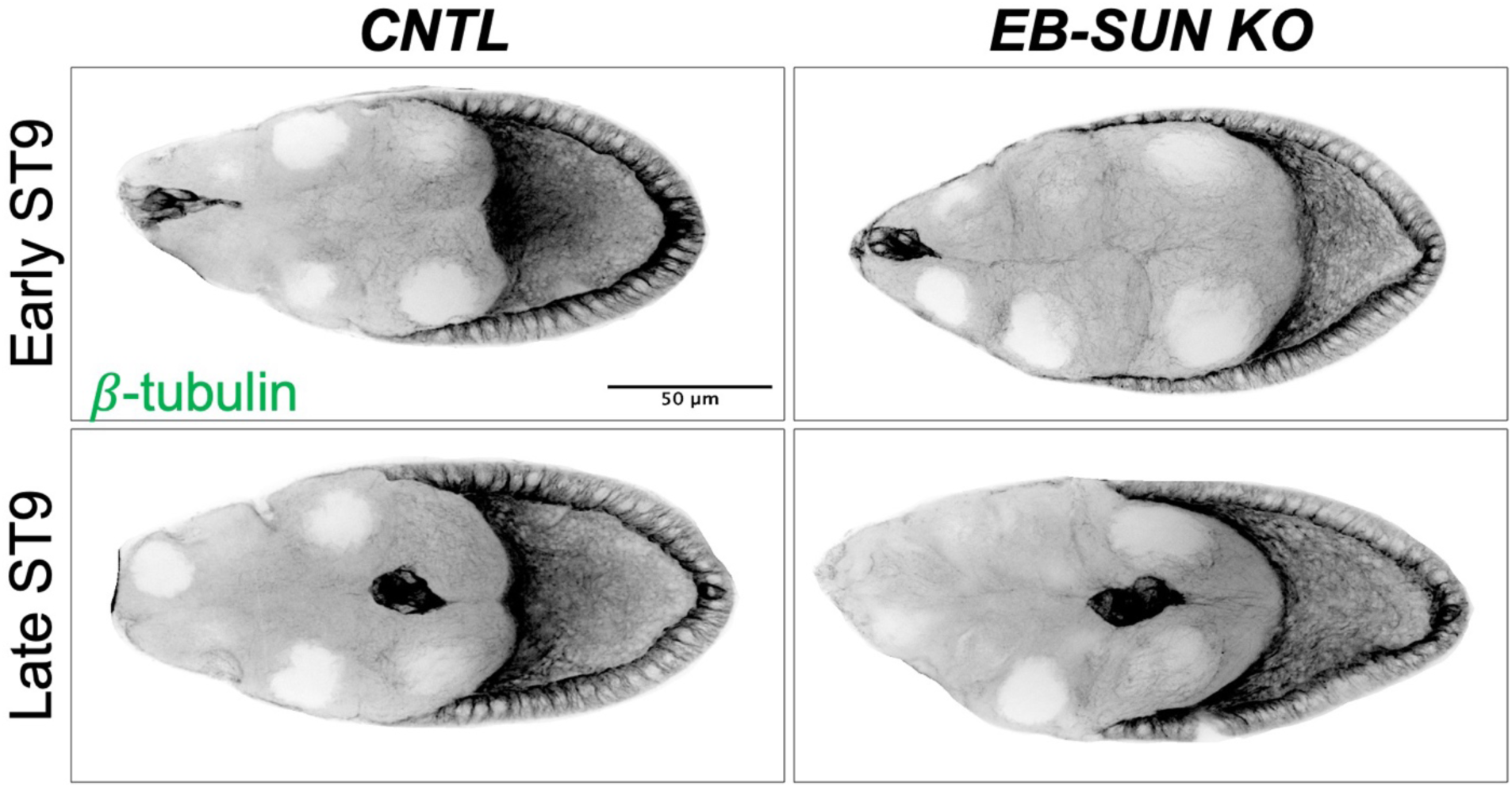
Microtubule staining of egg chambers at ST9. No significant difference in microtubule staining was observed between control and EB-SUN KO egg chambers.

**Figure S4.**
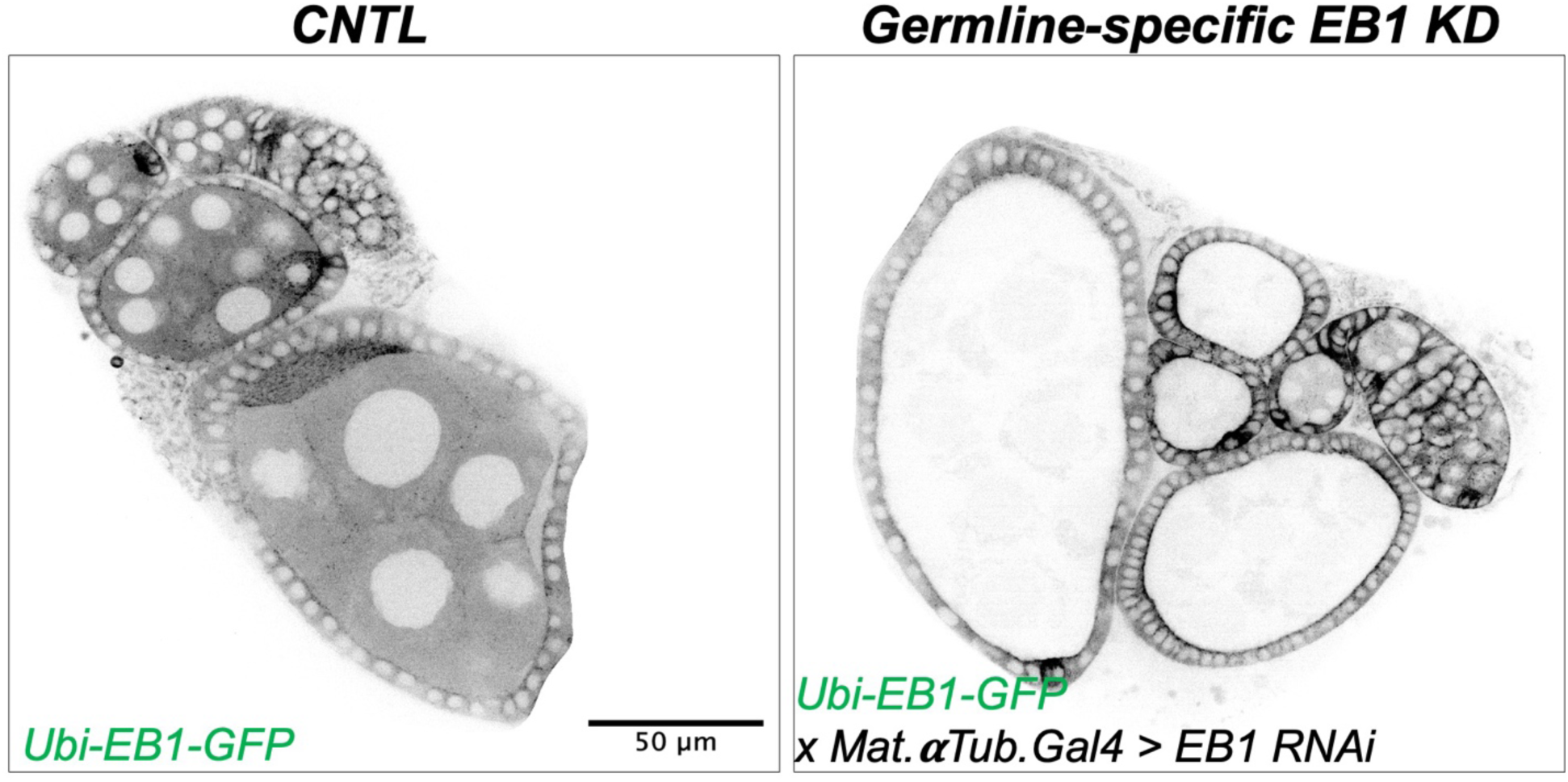
Germline-specific KD of EB1. To test EB1 RNAi efficiency, a fly line expressing GFP-tagged EB1 under a ubiquitin promoter (*ubi-EB1-GFP*) was crossed with an *EB1 RNAi* line driven by a germline-specific Gal4 (*Mat.αTub-Gal4^[V37]^*). As the germline-specific *Mat.αTub-Gal4^[V37]^* turns on between ST2 and ST3 after cell divisions, EB1 comets are completely abolished in germline cells (nurse cells and oocytes) within egg chambers past ST3, without affecting follicle cells. In control, ubi-EB1-GFP was crossed with *Mat.αTub-Gal4^[V37]^*alone. (See also Supplemental Movie S5)

**Figure S5.**
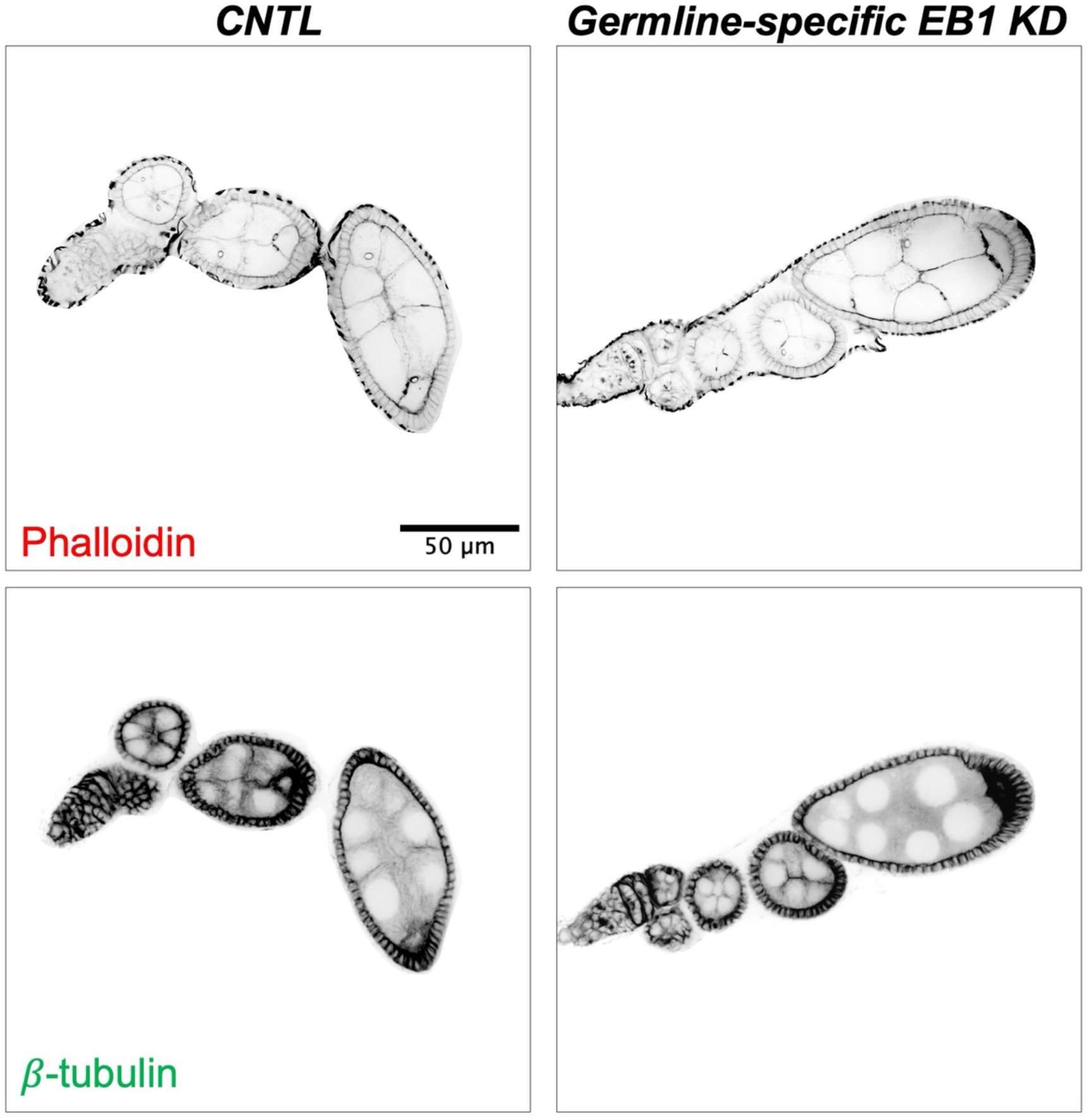
Germline-specific KD of EB1. EB1 was depleted by expressing EB1 RNAi by two Gal4 lines: *nos-Gal4^[vp16]^* which activates RNAi in primordial germ cells and a postmitotic germline-specific Gal4 (*Mat.αTub-Gal4^[V2H]^*), which turns on between ST2 and ST3 to ensure continuous depletion of EB1 throughout oogenesis. Overall, EB1 depletion in early cell divisions and oogenesis appears to be largely unaffected. Scale Bar, 50 µm

**Movie S1 (Related to Figure 1):** EB-SUN comets in *Drosophila* S2 cells. S2 cells were transiently transfected with a TagRFP-tagged EB-SUN construct. The movie shows EB-SUN localization at growing MT plus ends. Images were captured every second for 1 minute. The movie is displayed at 10 frames per second. Scale Bar, 10 µm.

**Movie S2 (Related to Figure 2):** EB-SUN and EB1 comets in *Drosophila* S2 cells. S2 cells were transiently transfected with a TagRFP-tagged EB-SUN and GFP-tagged EB1 constructs. The movie shows colocalization of EB-SUN with EB1 at growing MT plus ends. Images were captured every 2-second for 2 minutes. The movie is displayed at 10 frames per second. Scale Bar, 10 µm.

**Movie S3 (Related to Figure 4, B):** EB-SUN localization in the early oogenesis. *UAS-EB-SUN-1XTagRFP* expresses driven by an early germline driver, *nanos-Gal4^[VP16]^*. Images were captured every 2-second for 2 minutes. The movie is displayed at 10 frames per second. Scale Bar, 10 µm.

**Movie S4 (Related to Figure 4, C):** EB-SUN localization in the mid oogenesis. *UAS-EB-SUN-1XTagRFP* expresses driven by a germline-specific driver, *Mat.αTub-Gal4^[V37]^*. Images were captured every 2-second for 2 minutes. The movie is displayed at 10 frames per second. Scale Bar, 10 µm.

**Movie S5 (Related to Figure 6 and S5):** Germline-specific EB1 KD. EB1 comets are completely abolished in both nurse cells and oocytes by EB1 RNAi to confirm depletion of EB1. Images were captured every 2-second for 2 minutes. The movie is displayed at 10 frames per second. Scale Bar, 50 µm.

## Abbreviations

MT: Microtubule
KD: Knockdown
KO: Knockout
CH: Calponin Homology
MAP: Microtubule-associated protein
scRNA: single-cell RNA
ST: Stage

